# LACCASE is Necessary for Root Development in *Salvia miltiorrhiza*

**DOI:** 10.1101/2020.10.26.356097

**Authors:** Zheng Zhou, Qing Li, Yun Wang, Liang Xiao, Qitao Bu, Kai Hao, Meili Guo, Wansheng Chen, Lei Zhang

**Author notes:** These authors contributed equally to this article (Zheng Zhou, Qing Li). Author for correspondence: Meili Guo, Wansheng Chen, Lei Zhang 86+ 021-81871307.

## Abstract

Laccases are multicopper-containing glycoproteins related to monolignol oxidation and polymerization. These properties indicate that laccases are involved in the formation of important medicinal phenolic acid compounds in *Salvia miltiorrhiza* such as salvianolic acid B (SAB), which is used for cardiovascular disease treatment. To date, 29 laccases have been found in *S. miltiorrhiza*, some of which influence the synthesis of phenolic acids. Because of the functional redundancy of laccase genes, their roles in *S. miltiorrhiza* are poorly understood. In this study, the CRISPR/Cas9 system was first used for dual gene locus targeting in *S. miltiorrhiza* to knock out multiple laccase family genes. The development of the roots was retarded, and root microstructure was abnormal in laccase mutant lines. Additionally, the accumulation of phenolic acid compounds as well as lignin was dramatically reduced. This study suggests that SmLACs are necessary for root development and phenolic acid compound metabolism in *S. miltiorrhiza*.

## Introduction

*Salvia miltiorrhiza*, also known as Danshen, is a popular and effective traditional Chinese medicine used to treat cardio-cerebral vascular diseases(Guo *et al*., 2014). The key components of *S. miltiorrhiza* are phenolic acids such as salvianic acid (Danshensu), rosmarinic acid (RA), salvianolic acid A (SAA) and salvianolic acid B (SAB). Previous studies have shown that SAB, which accumulates at higher levels in roots than in other organs, may relieve ischemic cardiovascular diseases by promoting angiogenesis to improve microcirculation(Yang *et al*., 2016). Currently, the biosynthesis pathway of phenolic acids in *S. miltiorrhiza* has only been elucidated from phenylalanine and tyrosine to RA, and the biosynthesis of compounds with complicated structures, such as SAB, has yet to be revealed(Di P *et al*., 2013) (Supplementary Figure S1). To determine the synthetic metabolism of these phenolic acid compounds, the analysis and comparison of structural analogs to mine key enzymes is a good method.

Phenolic acids are lignin-derived products. Lignins are aromacic biopolymers derived from *p*-coumaryl, coniferyl and sinapyl alcohol (monolignol) oxidative coupling(Tang *et al*., 2019). These compounds deposit secondarily thickened cell walls, in which they play an essential roles in structural mechanical support, water transport and pathogen attack resistance(Ferrer *et al*., 2008; Burton *et al*., 2010). The polymerization of monolignols is facilitated by laccase proteins. Laccases are a multicopper oxidase family belonging to the benzenediol oxygen reductases (EC 1.10.3.2) and are also known as *p*-diphenol oxidases or urushiol oxidase(Festa *et al*., 2008; Wang *et al*., 2015). Laccases exist widely in nature and can be extracted from various species, such as bacteria, fungi, plants and mosses(Arregui *et al*., 2019). There are three conserved catalytic sites in laccase proteins, referred to as Type 1, Type 2 and Type 3 sites. In oxidization reactions, the substrate is first binds and is oxidized at the Type 1 site and its then transferred to a Type 2/Type 3 trinuclear copper cluster, with the release of an electron. An oxygen molecule accepts the electron, ultimately generating water(Wang *et al*., 2019). Because of the low substrate specificity of this enzyme family, most phenolic and nonphenolic molecules can be oxidized by laccases, and they are extensively applied in sewage treatment, lignin degradation and electricity-catalyzing reactions(Viswanath *et al*., 2014; Chandra & Chowdhary, 2015).

In planta, laccase was first identified from the Japanese lacquer tree, *Rhus vernicifera*, and its function was revealed in a series of studies. In *Arabidopsis thaliana*, 17 putative laccase genes have been identified from the genome, and AtLAC 4, AtLAC 17 and AtLAC 11 have been verified to be related to monolignol polymerization(Berthet *et al*., 2011; Zhao *et al*., 2013). *Oryza sativa* laccase may be involved in pesticide catabolism or detoxification(Huang *et al*., 2016). Regarding the influence of phenolic compounds, the overexpression of GaLAC1 from *Gossypium arboretum* in *A. thaliana* resulted in an increase in sinapic acid conversion into a monolactone-type dimer(Wang *et al*., 2004). In *S. miltiorrhiza*, after silencing a single SmLAC gene, the content of RA and SAB in transgenic hairy roots was reduced(Li *et al*., 2019).

Because of the functional redundancy of laccase genes, the decreasing the expression of a single has little influence on the plant phenotype. Therefore, it is necessary to silence or knock out multiple genes in the laccase family to comprehensively evaluate their roles. Due to the high homology of conserved domains in 29 laccase genes, we used the CRISPR/Cas9 system to edit conserved domains to target several laccase genes simultaneously. This is the first study to select multiple loci for editing in *S. miltiorrhiza*. The editing targets were selected among 29 SmLACs based on bioinformatics analysis and nucleotide and amino acid sequence alignments. Then, the targets were introduced into a binary vector to construct a single- or double-locus editing vector, which was transformed into hairy roots via the *Agrobacterium rhizogenes* method. The results showed that more than 20 SmLACs were knocked out with this system. Quantitative real-time PCR and RNA-Seq experiments were performed to better understand the impact of SmLAC suppression, and the results demonstrated that the expression of target genes was decreased. The development of hairy roots was significantly suppressed in the gene-edited lines, and the RA and SAB contents decreased dramatically. The examination of cross-sections of laccase mutant hairy roots showed that root development was abnormal and that the xylem cells in the edited samples appeared larger and looser than those in the wild type. Furthermore, lignin was nearly undetectable. In conclusion, our study showed that SmLACs play an important role in lignin formation and are necessary for root development in *S. miltiorrhiza*.

## Methods and materials

### Conserved domain alignment and sgRNA design

Conserved motifs in the complete amino acid sequences of SmLACs were identified using Multiple EM software for Motif Elicitation with the setting of a maximum of 10 motifs(Bailey *et al*., 2006). Alignments of SmLAC gene amino acid and nucleotide sequences were performed with MEGA 5.05 software with p-distance settings. The nucleotide sequences of highly conserved domains were selected and used in the online CRISPR/Cas9 design tool CHOPCHOP (http://chopchop.cbu.uib.no/) to search for potential gene editing sites. Every possible editing target followed by 5’-NGG was evaluated and ranked considering the off-target possibilities and editing location. Sequences with high marks are ideal targets for editing.

### Vector construction

Pairs of complementary oligos from selected sgRNAs were synthesized and annealed to generate dimers. For single-target-editing vector construction, the sgRNA1 dimer was directly cloned into the CRISPR/Cas9 system by B*bs*I digestion. For dual-targets-editing vector construction, the sgRNA2 dimer was first cloned into the CRISPR/Cas9 system, and the cassette including AtU6 and sgRNA2 was then amplified with *Kpn*I and E*coR*I restriction sites and subcloned into the single-target-editing vector. Finally, two CRISPR/Cas9 vectors were subcloned into the linearized pCambia1300 (Cambia, Canberra, Australia) plant expression vector by double restriction enzyme digestion and recombination.

### Plant materials

The *S. miltiorrhiza* cultivar (RA content, 3.5-3.8% DW, LAB content, 2.8-3.0 % DW) used in this study was selected and developed over several years at the Second Military Medical University. The procedures for obtaining aseptic seedlings, *Agrobacterium-* mediated transformation and hairy root culture performed as previously described(Zhou *et al*., 2018).

### Transgenic hairy root identification and target gene mutation detection

DNA was extracted from the obtained hairy roots via the CTAB method, and two pairs of primers were used to detect the *rolB* and *cas9* genes from the C58C1 strain as well as binary vector by PCR amplification. The fragments surrounding LAC10, LAC11, LAC13, LAC16, LAC21, LAC27 and LAC28 were amplified by PCR using the primers listed in Supplementary Table S1. Gene editing induced mutations were detected by aligning the sequencing chromatograms of these PCR products with those of the WT controls. To identify the genotypes of the mutations, PCR products were cloned into a blunt zero vector (Transgen Biotech, Beijing) for sequencing.

### Relative expression analysis via quantitative real-time PCR

Total RNA was extracted from samples using the EasyPure Plant RNA Kit (Transgen Biotech, China) and then reverse transcribed to produce cDNA using a TransScript First-Strand cDNA Synthesis SuperMix Kit (TransGen Biotech). Quantitative real-time polymerase chain reaction (qRT-PCR) assays were performed as reported previously(Zhou *et al*., 2018). All of the primers used for qRT-PCR are listed in Table S1.

### RNA-seq analysis

Total RNA was extracted from the hair roots of *Cas1-3, CAS12-5* and WT using the TRIzol reagent (TransGen Biotech, China). RNA quality was examined on a Bioanalyzer 2100 spectrophotometer (Agilent), and the RNA was stored at −80°C. RNA samples with an RIN >8.0 were considered suitable for subsequent experiments. cDNA libraries were prepared using the NEBNext™ Ultra Directional RNA Library Prep Kit (NEB, USA), the NEBNext Poly(A) mRNA Magnetic Isolation Module, and NEBNext Multiplex Oligos according to the manufacturer’s instructions. The products were purified and enriched by PCR to produce the final cDNA libraries and were quantified on an Agilent 2100 spectrophotometer. The tagged cDNA libraries were pooled in equal ratios and used for 150 bp paired-end sequencing in a single lane on the Illumina HiSeqXTen platform. The fragment counts of each gene were normalized via the fragments per kb per million (FPKM) method.

### Histochemical staining of hairy roots

Histochemical staining was performed on sections cut from the maturation zone. The hairy roots were embedded in 7% agarose before being transversely sectioned at a thickness of 50 μm using a vibratome (Leica VT1000S, Leica, Germany). The safranin O-green staining method was used to detect the deposition and composition of lignin(Bouvier *et al*., 2013). All sections were observed under an Olympus BX43 microscope (Japan).

### RA and SAB content determination

Hairy roots of different lines were collected and dried at 40°C, and 0.1 g samples were then used to measure RA and SAB contents by high-performance liquid chromatography-tandem mass spectrometry as reported previously (Agilent 1200, USA)(Zhou *et al*., 2018).

## Results

### Conserved domain alignment of SmLACs and editing site selection

The complete open reading frames (ORFs) of 29 SmLACs were obtained from a previous study, and each of them was analyzed for conserved domains with the BLAST tool(Li *et al*., 2019). Although the lengths of the sequences were different, 3 conserved copper atom binding sites commonly existed among these SmLAC proteins.

Two conserved domain sequences were selected as target loci for editing, and the online web tool CHOPCHOP was used to search for suitable sgRNA sequences for the CRISPR/Cas9 editing system. Potential editing sites were ranked according to software scores based on several factors, such as self-complementarity and possible editing efficiency. Finally, two 20 bp gene editing sites were identified: sgRNA1: GCCAACTATATATGCTCGAA, corresponding to a conserved Type 1 catalytic site, and sgRNA2: CAATGCATAAACCACACCCC, corresponding to a conserved Type 2 domain (Figure 1A, B).

**Figure 1.**
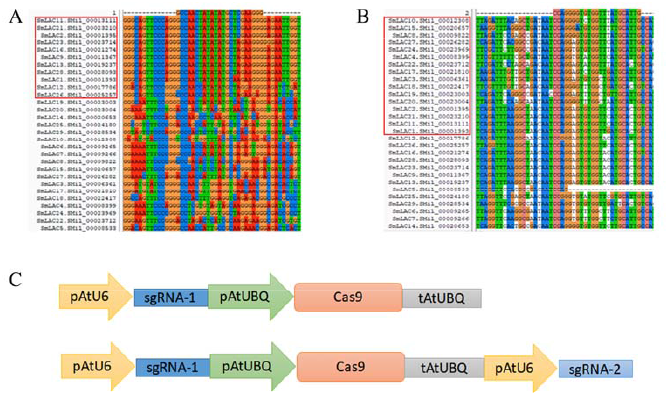
Overview of the CRISPR/Cas9 system for SmLAC disruption. (A and B) Partial sequence alignment of the first and second conserved copper domains among 29 SmLACs. The target sequence listed above and PAM sequence (GGG) are indicated with blue bars. The highly conserved regions from different SmLACs aligned with the editing targets are indicated (red frame). (C) Schematic view of two CRISPR/Cas9 knockout vectors used in this study. The cas9 protein was driven by the ubiquitin promoter, and the sgRNA was driven by the U6 promoter of Arabidopsis. The *Cas1* vector carried only one AtU6-sgRNA cassette, and the *Cas12* vector carried two AtU6-sgRNA cassettes to induce double mutations in SmLACs.

The sgRNA1 sequence existed in 11 SmLACs (LAC1, LAC2, LAC9, LAC11, LAC12, LAC13, LAC16, LAC21, LAC23, LAC26, LAC28), and the sgRNA2 sequence existed in 16 SmLACs (LAC1, LAC2, LAC3, LAC4, LAC8, LAC10, LAC11, LAC15, LAC17, LAC 18, LAC 19, LAC20, LAC21, LAC22, LAC24, LAC27). Both the sgRNA1 and sgRNA2 sequences existed in 4 genes (LAC1, LAC2, LAC11, LAC21).

### CRISPR/Cas9 expression vector construction

The CRISPR/Cas9 system was provided by Jian-Kang Zhu’s laboratory at the Shanghai Center for Plant Stress Biology, Chinese Academy of Sciences. In the vector, the sgRNA was driven by the U6 promoter, and the Cas9 protein was driven by the ubiquitin promoter from *A. thaliana*. To determine the editing efficiency for multiple genes in *S. miltiorrhiza*, we constructed two vectors. To construct the single-locus knockout vector, the 20 bp sgRNA1 sequence was inserted between the AtU6 and ubiquitin promoters. To generate the dual-site editing vector, a new cassette including the extra AtU6 promoter and sgRNA2 was introduced into the single-locus knockout vector (Figure 1C). The whole expression cassette was subcloned into the pCAMBIA1300 plant expression vector for *A. tumefaciens-mediated* transformation. The vector carrying sgRNA1 was referred to as *Cas1*, and the vector carrying both sgRNA1 and sgRNA2 was referred to as *Cas12*. The empty pCambia1300 vector was used as a control.

### Transgenic hairy root identification and SmLAC expression analysis

Sixteen strains of transgenic hairy roots were obtained in both the *Cas1* and *Cas12* lines, and PCR amplification was used to identify the positive transgenic hairy roots. The results showed that the CRISPR/Cas9 vector was successfully transformed into 15 transgenic hairy roots in *Cas1* lines and 14 transgenic hairy roots in *Cas12* lines (Figure 2G and H). Four transgenic hairy root strains from both lines were used in subsequent studies.

**Figure 2.**
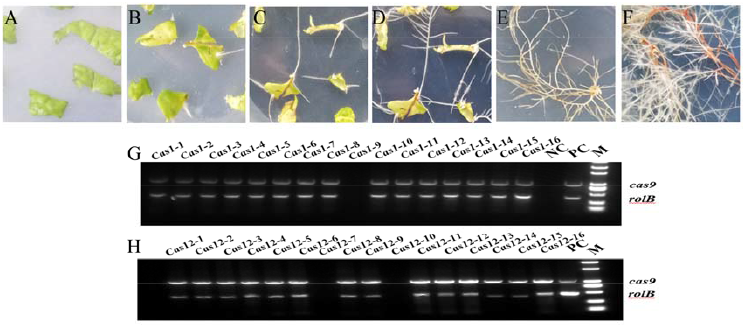
Cultivation transgenic hairy roots and identification of positive lines. (A) The leaf discs are spread on MS medium after *Agrobacterium-mediated* infection and undergo (B) callus formation, (C) hairy root growth, (D) bifurcation, (E) strain separation and (F) growth cultivation to obtain mature roots for analysis. (G and H) Identification of the positive transgenic hairy roots using PCR amplification. The *rolB* genes from the *A. rhizogenes* C58C1 strain and the cas9 gene from the constructed vector were amplified with two pairs of primers. The empty vector line was used as a positive control (PC), and wild-type hairy roots were used as a negative control (NC).

Five laccase genes from editing domains were selected to determine the effect of the CRISPR/Cas9 system by quantitative real-time PCR. The results showed that the expression levels of the target SmLACs decreased dramatically after editing (Figure 3A and B). To better understand the impact of laccase editing, *Cas1-3, Cas12-5* and wild-type (WT) plants were evaluated via RNA-Seq experiments. A total of 57,987,044, 57,609,184 and 60,855,444 sequencing reads were successfully produced for WT, *Cas1-3* and *Cas12-5*, respectively. Moreover, compared with the WT line, 3108 and 2666 differentially expressed genes were identified from the *Cas1-3* and *Cas12-5* strains with an FDR < 0.05 and log2|FC|>1.

**Figure 3.**
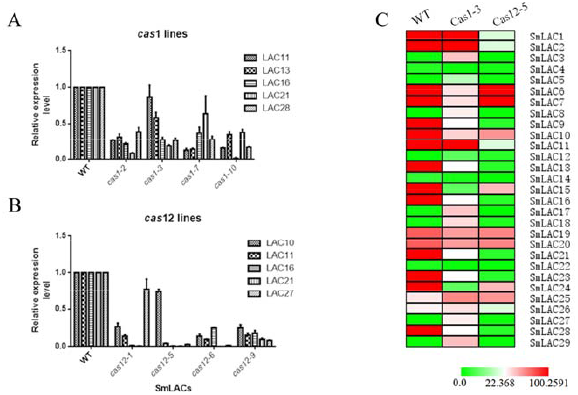
Expression profiles of SmLACs in transgenic hairy root lines. (A) Expression levels of five laccase genes (SmLAC11, SmLAC13, SmLAC16, SmLAC21, SmLAC28) in *Cas1* lines by quantitative real-time PCR determination. (B) Expression levels of five laccase genes (SmLAC10, SmLAC11, SmLAC16, SmLAC21, SmLAC27) in *Cas12* lines. (C) Heat maps of 29 SmLACs in the wild-type, *cas1-3* and *cas12-5* lines. Rows represent differentially expressed laccase genes, and columns represent group comparisons. Red and green boxes represent lower and higher gene expression levels, respectively. The depth of color demonstrates the absolute value of the magnitude of the base-2 logarithmic fold change expression ratio.

We identified a total of 29 SmLAC genes on the basis of transcriptome statistics and used a heat map to demonstrate the expression differences clearly (Figure 3C). The results showed that the expression levels of the edited laccase genes were decreased, and this result was in accordance with qRT-PCR detection. In the *Cas1-3* strain, 6 SmLACs (LAC9, LAC13, LAC16, LAC21, LAC23 and LAC28) showed dramatic decreases after gene editing. In addition, the expression levels of 9 SmLACs in the *Cas12-5* strain were reduced, and 4 SmLACs (LAC1, LAC2, LAC11, and LAC21) presented markedly lower levels than were found in *Cas1-3*. These results indicate that the expression levels of most SmLACs were markedly decreased in the transgenic edited lines.

To detect the mutations in the *Cas1-3* and *Cas12-5* strains, selected SmLAC regions (approximately 500 bp) containing the target sequences were amplified by using specific primers. The purified PCR products were sequenced directly, and the corresponding chromatograms were decoded using the Degenerate Sequence Decoding method. *S. miltiorrhiza* is a diploid plant, and the CRISPR/Cas9 system can lead to two mutation types in the target genes. A simplex peak suggests that a mutation appears on both strands and can be designated a homozygous mutation. An overlapping peak indicates that a transgenic hairy root is heterozygous and contains a mutation that appears on a single strand or that different mutations occur on the two DNA strands. All the PCR products from heterozygotes were cloned into a sequencing vector to identify mutations on different strands.

Among the successfully target-edited lines, most of the mutation events consisted of 1 or 2 nucleotide deletions or insertions. However, the loss of large fragments was also detected, as found in *Cas12-5-LAC*21 (Supplementary Figure S2). In most cases, the insertion or deletion of nucleotides influenced SmLAC translation, resulting in open reading frame shifts or early termination of target genes. The results indicated that it is feasible to knock out multiple targets with a single binary vector harboring several sgRNAs for gene editing in *S. miltiorrhiza*.

### Phenotypic analysis of CRISPR/Cas9-generated transgenic hairy roots

To evaluate the phenotype of the transgenic hairy roots, the growth status, microstructure and metabolite content of the WT and edited lines were collected. Compared with the thriving growth of the control lines, the SmLAC mutant lines produced hairy roots showing retardation of root growth with the emergence of sparse root hairs (Figure 4A-F). The roots of the *Cas1* lines were thin, and the hairy roots of almost all of these lines showed no or only one to two lateral roots distributed along the axial root (Figure 4G-I).

**Figure 4.**
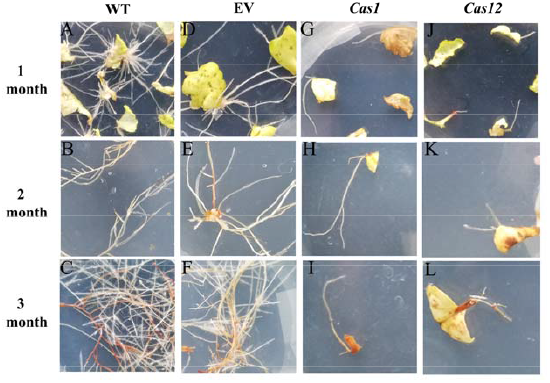
Phenotype of genome-edited plants. Hairy root growing state at 1 month, 2 months and 3 months after *Agrobacterium-mediated* infection of (A-C) wild-type lines, (D-F) empty vector lines, (G-I) *Cas*1 lines and (J-L) *Cas*12 lines.

Moreover, the root growth of the *Cas12* lines was retarded and stopped at the elongation stage (Figure 4J-L). Additionally, the number of hairy roots extending from the callus was strikingly different among the control and gene-edited lines. Compared with the control lines, the SmLAC mutant lines developed fewer regenerated roots from calli.

To investigate the root microstructure and changes in lignin after gene editing, transverse sections of the hairy roots of *Cas1-3, Cas12-5* and WT plants were studied, and safranin O-green staining was performed. This staining results in the development of a red stain in the presence of lignified cell walls. It was found that lignin accumulation was rarely observable in the mutant lines compared with the WT lines. Moreover, even the normally strongly lignified xylem showed no obvious lignin staining in the SmLAC-edited lines. Higher magnification observations of root transverse sections suggested that very few vessels were lignified in the mutants, whereas lignification was observed in most of the xylem vessels in the WT lines. Additionally, the SmLAC-edited lines showed cellular structural abnormalities, reflected in differences in root cell shape and organization. The arrangement of the xylem and phloem of the SmLAC-edited lines was completely different from that in the controls. In the WT and empty vector lines, the xylem vessels of the roots were radially arranged, while the phloem was alternately arranged in concentric rings (Figure 5A-F). However, as observed in the *Cas1* strains, the xylem was arranged sparsely, and some cells in the phloem appeared to be collapsed (Figure 5G-I). According to the *Cas12* strains, disorganized vascular patterning involving xylem cells developed in regions where phloem cells were distinctly enlarged and irregular (Figure 5J-L).

**Figure 5.**
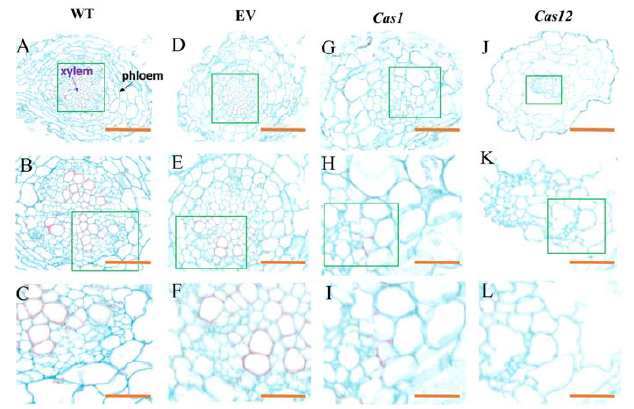
Phenotype of root microstructure in laccase mutant observed in cross-sections. The microstructure of (A-C) wild-type lines, (D-F) empty vector lines, (G-I) cas1 lines and (J-L) cas12 lines. The purple arrow indicates the xylem, and the black arrow indicates the phloem. The green rectangles represent enlarged regions. Bars, A, D, G, J, 100μm, B, E, H, K, 50 μm, C, F, I, L, 20 μm.

To explore the influence of the SmLAC mutations on phenolic acid synthesis, the identification of RA and SAB in regenerated hairy root extracts was performed via the HPLC/MS method; retention times were assessed in comparison with authentic standards as well as by searching data in the literature. The results showed that RA and SAB contents were decreased in all edited lines, in which they were approximately 1-3 times lower than in WT lines. Furthermore, the metabolite content of the *Cas12* lines was dramatically reduced compared with that in *Cas1* lines (Figure 6).

**Figure 6.**
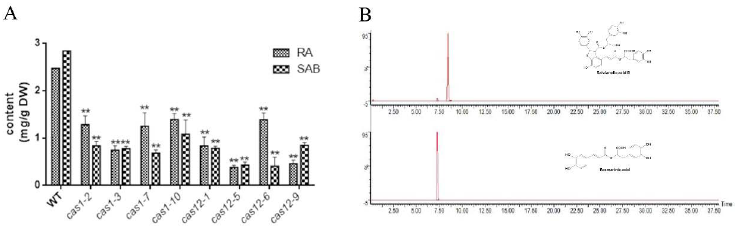
Determination of the contents of RA and SAB in the hairy roots of the experimental lines. (A) The contents of RA and SAB in transgenic hairy roots. (B) Chromatograms of the hairy root extract from the transgenic lines relative to an RA standard and an SAB standard. All data presented here are means ± S.D. of three biological replicates, with error bars indicating standard deviations. Statistical significance was determined by the Student’s t test (* P < 0.05).

## Discussion

Laccases have been reported to play roles in processes such as the polymerization of monolignol, iron metabolism, and responses to environmental stresses(Hoopes & Dean, 2004; Shen *et al*., 2013). In contrast, although three laccase genes in *S. miltiorrhiza* (SmLAC7, SmLAC20 and SmLAC28) have been reported to influence phenolic acid synthesis, the roles of SmLACs in physiological processes remain less clear. In this study, we first report the roles of SmLACs in the alteration of lignin synthesis, which is essential for root development and phenolic acid ingredient metabolism. The retardation of hairy root growth and lateral root formation caused by CRISPR/Cas9 editing may result from the disturbance of diverse physiological and biochemical processes. The significant microstructural changes observed in transverse sections from SmLAC-edited lines also suggested that laccases play a key role in root development. In addition, the contents of RA and SAB were decreased dramatically, suggesting that SmLAC genes are involved in phenolic acid biosynthesis.

Phenolic acids can be derived from the processing of lignin, and in *S. miltiorrhiza*, the biosynthesis of phenolic acids starts with L-phenylalanine and L-tyrosine, which are separately transformed into the intermediates 4-coumaroyl CoA and 3,4-dihydroxyphenyllactic acid. Rosmarinic acid synthase and cytochrome P450-dependent monooxygenase catalyze two precursors to synthesize RA. Lactase has been proposed as a key enzyme involved in the synthetic process of RA to SAB conversion (Supplementary Figure S1)(Di P *et al*., 2013). Therefore, it is essential to understand this enzyme family. Here, we studied laccase function from a comprehensive perspective, focusing not only on phenolic acid metabolism but also on the alteration of physiological processes in *S. miltiorrhiza*.

Despite the high degree of redundancy of laccase genes in *S. miltiorrhiza*, it is difficult to identify lignin-specific enzymes. It is time consuming to generate multiple mutants to study the function of multiple genes. In gene family function studies in particular, the function of a single gene might be redundant, and multiple genes should be knocked out to explore their phenotypes and roles. Therefore, this study provides new gene editing methods to study this gene family in *S. miltiorrhiza*. In previous studies, the CRISPR/Cas9 system was reported to target a single gene locus at a time, and this study is the first to perform dual-locus editing(Li *et al*., 2017; Zhou *et al*., 2018; Deng *et al*., 2020). This work will facilitate the application to a new strategy for functional studies, targeting multiple genes and gene families in *S. miltiorrhiza*.

In addition to laccase, other classes of oxidative enzymes, such peroxidases and polyphenol oxidases, are also speculated to participate in lignin polymer formation(Zamocky *et al*., 2015). Peroxidases and laccases that are potentially involved in the oxidation of lignin precursors in plant cell walls are encoded by multigene families(Ruiz-Duenas *et al*., 2009). The peroxidase-catalase superfamily may be further subdivided into three classes. Class I involves intracellular peroxidases, and class II peroxidases are extracellular fungal peroxidases involved in lignin degradation, including lignin peroxidase (LiP), manganese peroxidase (MnP), and versatile peroxidase (VP). Class III includes peroxidases secreted by plants, such as horseradish peroxidase, which has been implicated in cell wall biosynthesis, indole-3-acetic acid catabolism and the oxidation of poisonous compounds. Among these three subfamilies, Class II has attracted the most attention because of the catalytic activity of lignin. While MnP oxidizes phenolic structures of lignin and LiP targets nonphenolic components, VP has the ability to oxidize both phenolic and nonphenolic structures. Unlike laccase, which uses oxygen to oxidize its substrates, peroxidases consume hydrogen peroxide to perform the reaction(Falade *et al*., 2017). In *S. miltiorrhiza*, 73 peroxidases have been identified, and the function of these peroxidases still needs to be explored.

## Acknowledgements

This work was funded by National Natural Science Foundation of China (Grant nos., 31770329, 32070327), National Key R&D Program of China (2019YFC1711100).

## Author contributions

Z.Z., M.G., W.C. and L.Z. conceived and designed the entire research plans; Z.Z., Q. L. and L.X., performed most of the experiments; Q. L., Y. W., and Q.B., K. H., provided technical assistance to Z.Z.; Z.Z. and Z. L. wrote the manuscript; and M. G. and W.C. helped with the organization and editing.

